# Caspase-1 and -11 defend against environmental but not host-adapted pathogens in genetically complex mice

**DOI:** 10.64898/2026.10.30.763184

**Authors:** Lupeng Li, Benjamin D. McGlaughon, Heather N. Larson, Beverly H. Koller, Edward A. Miao

## Abstract

Caspase-1 and -11 are inflammatory caspases that trigger pyroptosis, yet in most infection models they confer only a modest survival benefit. We have proposed that caspase-1 and -11 evolved primarily to defend against ubiquitous environmental microbes that have not been under selective pressure to evade them, whereas host-adapted pathogens have evolved evasion strategies. Testing this idea is complicated because the inbred mouse strains used in most infection studies carry mutations in innate immune genes: i.e., C57BL/6 (B6) mice carry a loss-of-function *Nramp1* allele, and 129 mice lack functional caspase-11. Here we generated B6 × 129 F1 hybrid mice, which are gene-sufficient at both *Nramp1* and *Casp11*, and used them to measure the fold change in lethal dose (ΔLD_100_) caused by caspase-1 and -11 deficiency for four pathogens on the B6, 129, and F1 backgrounds. Caspase-1 and -11 conferred little or no survival benefit against the host-adapted pathogens *Listeria monocytogenes* and *Salmonella* Typhimurium on any of the three backgrounds, apart from a modest benefit at the lowest *S*. Typhimurium inoculum in F1 mice. In contrast, caspase-1 and -11 were required to survive infection with the environmental pathogens *Burkholderia thailandensis* and *Chromobacterium violaceum*, where caspase-1 and -11 deficiency lowered the lethal dose by up to six orders of magnitude. The failure of caspase-1 and -11 against host-adapted pathogens therefore persists on a genetically robust hybrid background, supporting the view that a major evolutionary role of caspase-1 and -11 is defense against environmental microbes that have not been selected to evade them.

**Importance:** Cells can sense bacteria growing inside them and destroy themselves, killing the invader along with the cell. Two enzymes, caspase-1 and caspase-11, carry this out, yet mice lacking both die from host-adapted bacteria such as *Salmonella* and *Listeria* almost as readily as normal mice. We proposed that these enzymes instead evolved to stop the ubiquitous bacteria of soil and water, microbes that have never faced selective pressure to hide from them. Testing this has been difficult because the standard inbred strains each carry defective immune genes. We crossed two strains to produce hybrid mice with a complete set, then measured the dose of bacteria needed to kill them. Caspase-1 and caspase-11 gave almost no protection against *Salmonella* or *Listeria*, but raised the lethal dose of the environmental bacteria *Burkholderia thailandensis* and *Chromobacterium violaceum* by up to a millionfold. This defense therefore appears aimed at environmental microbes.

## Introduction

The Red Queen’s race holds that a species must evolve continuously merely to hold its ground, because the organisms it competes with are evolving in step with it^1^. Host–pathogen interactions are a canonical arena for this race. Hosts elaborate immune defenses, and host-adapted pathogens, which can be personified as Red Queens, answer them with virulence factors that subvert those same defenses. Each competitive advancement prompts a reciprocal response, ensuring that neither adversary maintains a permanent advantage.

Pyroptosis is a lytic form of programmed cell death that releases inflammatory cellular contents and is executed by the inflammatory caspases, caspase-1 and caspase-11. Caspase-1 is activated within inflammasomes, cytosolic multiprotein complexes that detect microbial contamination and danger signals^2^. Caspase-11 is instead activated by directly binding LPS that has gained access to the cytosol, without requiring a sensor complex. Both caspases cleave gasdermin D, which forms the membrane pores that execute pyroptosis. Therefore, caspase-1 and -11 should help to control intracellular pathogenic infections by removing replicative niches, recruiting immune cells, and inducing inflammation^3,4^. Most published studies on the role of pyroptosis in defense against infection have focused on host-adapted pathogens. The general conclusion is that mice lacking caspase-1 and -11 have increased susceptibility to infection.

*Salmonella enterica* serovar Typhimurium, a host-adapted pathogen, has been extensively studied in caspase-1/11-deficient mice^5–12^. The NLRC4 inflammasome can be activated in primary macrophages by purified flagellin, rod, or needle proteins of *S*. Typhimurium^13^. Rod and needle are protein components of the *S*. Typhimurium type III secretion system (T3SS) apparatus encoded by *Salmonella* pathogenicity island-1 (SPI1)^14^. However, during systemic infection, *S*. Typhimurium evades NLRC4 inflammasome detection by repressing flagellin expression, and expressing the SPI2-encoded T3SS rod protein SsaI, which is undetectable by the NLRC4 inflammasome^15^. Although the SPI2 needle protein, SsaG, can be detected by the NAIP-NLRC4 inflammasome when artificially delivered to the cytosol in mouse macrophages, during infection with live *S*. Typhimurium, this agonist fails to trigger the downstream pyroptotic pathway perhaps due to extremely low expression levels of the protein^16^. Furthermore, *S*. Typhimurium primarily resides and replicates in the vacuolar space of host cells, preventing its LPS from being detected by caspase-11^15^. A subpopulation does escape into the cytosol, particularly in intestinal epithelial cells, where it can be detected^17^; however, this population is a minority during systemic infection. The caspase-1 and -11 evasion strategy is important for the virulence of *S*. Typhimurium, as re-expression of flagellin or SPI1 rod protein causes attenuation in the engineered *S*. Typhimurium strain in WT mice; in contrast, caspase-1/11-deficient mice remain fully susceptible to the engineered strains^8,18^. While caspase-1/11-deficient mice had increased *S*. Typhimurium burdens compared to WT mice, ultimately, both WT and caspase-1/11-deficient mice succumbed to infection; therefore, the fold increase in lethal dose (ΔLD_100_) between WT and caspase-1/11-deficient mice remained the same (Table 1 summarizes the previously published results). In other words, caspase-1 and -11 in WT mice fail to prevent death during *S*. Typhimurium infection. A major reason is that *S*. Typhimurium, as a successful host-adapted pathogen, evades caspase-1 and -11 detection and thereby renders caspase-1 and -11 mostly ineffective in preventing death^8,18,19^.

**Table 1.** Fold increase in LD_100_ (ΔLD_100_) in caspase-1/11-deficient mice. Calculated as described^15^.

| **Pathogen** | **ΔLD<sub>100</sub>** | **Ref.** |
|---|---|---|
| *Burkholderia thailandensis* | 6,000,000 | 28,29,60,69 |
| *Chromobacterium violaceum* | 100,000 | 29 |
| *Listeria monocytogenes* | >2 | 44 |
| *Salmonella* Typhimurium | 1 | 5,6,9,70–73 |
| 31 other examples | >8 | 15,26 |

Another host-adapted pathogen, *Listeria monocytogenes*, produces abundant flagellin at 22 °C but represses it at 37 °C, which is considered a virulence strategy^20^. In fact, *L. monocytogenes* engineered to express flagellin at 37 °C was highly attenuated in WT mice but remained virulent in *Nlrc4*^−/−^ mice^21,22^. Other host-adapted pathogens actively inhibit inflammasomes. For example, *Yersinia pseudotuberculosis* delivers the effector YopM, which recruits the kinases RSK1 and PKN to phosphorylate pyrin and thereby hold the pyrin inflammasome in an inactive state^23–25^. In two previous review articles, we estimated the ΔLD_100_ between WT and caspase-1/11-deficient mice across published studies of host-adapted pathogens, and nearly all infection models had ΔLD_100_ of less than 8 (Table 1), leading us to question the evolutionary importance of caspase-1 and -11 in defense against vertebrate-adapted pathogens that evade caspase-1 and -11 detection^15,26^.

Since the evasion strategies of host-adapted pathogens make caspase-1 and -11 inefficient at defending against infection, are there any pathogens that fail to evade caspase-1 and -11? Our laboratory has identified two environmental pathogens, *Burkholderia thailandensis* and *Chromobacterium violaceum*, which are successfully detected and cleared by caspase-1 and - 11. *B. thailandensis* is a Gram-negative bacterium encoding a T3SS related to *Shigella*^27^. Rod/needle proteins and LPS of *B. thailandensis* are detected by NLRC4 and caspase-11, respectively, leading to activation of caspase-1 and -11^28^. WT mice survived *B. thailandensis* infection with 6 × 10^7^ CFU and succumbed only to an extremely high dose of 6 × 10^8^ CFU infection, doses where even nonpathogenic, laboratory-adapted *E. coli* strains are lethal. In contrast, caspase-1/11-deficient mice died when infected by even 10^2^ CFU. Therefore, caspase- 1 and -11 changed the lethal dose by 6,000,000-fold. *C. violaceum* is another Gram-negative bacterium that encodes a T3SS related to *Salmonella*. The *C. violaceum* T3SS apparatus is detected by the NLRC4 inflammasome to facilitate bacterial clearance in vivo^29^. WT mice survived infection with 10^6^ CFU of *C. violaceum* but died after a very high dose of 10^7^ CFU infection, whereas caspase-1/11-deficient mice died after 10^2^ CFU infection. We estimated the ΔLD_100_ of *C. violaceum* between WT and caspase-1/11-deficient mice to be 100,000-fold. Therefore, compared to host-adapted pathogens, caspase-1 and -11 are extremely efficient at clearing specific environmental pathogens.

The striking ΔLD_100_ differences between host-adapted pathogens and environmental pathogens led us to propose that the primary physiological and evolutionary role of caspase-1 and -11 is to defend against pathogens that typically infect hosts lower on the evolutionary tree where caspase-1 and -11 are absent. We therefore proposed that caspase-1 and -11 achieved a near-permanent victory over environmental pathogens by forming “Red Pawn barriers” that prevent environmental pathogens from evolving in the Red Queen’s race. We can also call environmental pathogens “Red Pawn pathogens”. In contrast, host-adapted pathogens (Red Queen pathogens) have evolved evasion strategies to prevent detection by caspase-1 and -11. Caspase-1 and -11 thus act as a Red Pawn barrier: a defense that the affected pathogens fail to counter, demoting them out of the Red Queen’s race rather than driving another round of it (Miao, 2026, see attached manuscript that is under consideration at *Trends in Micro* for a detailed description). We define Red Pawns as ubiquitous environmental microbes that carry virulence traits (e.g., T3SS) but are readily cleared by the innate immunity of non-hosts (e.g., humans), because they are not in constant evolution with those non-hosts.

Although the ΔLD_100_ between host-adapted pathogens and environmental pathogens (Table 1) supports the Red Pawn barrier concept, two caveats of the prior studies prevent us from fully accepting it. Firstly, most published studies used one dose of infection rather than multiple doses; thus, the ΔLD_100_ calculated in Table 1 may be underestimated. Secondly, most studies used C57BL/6 (B6) mice to assess the function of caspase-1 and -11. However, B6 mice carry mutations in several innate immune genes. Most important for intracellular bacterial infection, B6 mice carry a loss-of-function allele of natural resistance-associated macrophage protein 1 (*Nramp1*), a gene that mediates protection against *S*. Typhimurium and *L. monocytogenes*^5,30^. Therefore, the susceptibility of *Casp11*^−/−^ mice to *B. thailandensis* infection may actually represent a dual gene deficiency where only *Nramp1*^mt/mt^*Casp11*^−/−^ mice are susceptible to *B. thailandensis*. Another commonly used inbred strain, 129 mice, lacks functional caspase-11 due to a 5-bp deletion^31^, making this strain susceptible to *B. thailandensis* infection^28^. Of note, 129 mice have a normal *Nramp1* gene, and B6 mice have an intact *Casp11* gene.

Since inbred mice may carry additional mutations in immune genes that have not yet been discovered, we decided to use F1 hybrid mice by crossing B6 and 129 mice (Figure 1A) to assess the importance of caspase-1 and -11 in defense against infection. F1 hybrid mice carry a complex assortment of SNPs and are genetically reproducible^32^. Moreover, the control F1 WT mice that we generated by crossing B6 WT mice with 129 WT mice are gene-sufficient at *Nramp1* and *Casp11*. As is typical for almost all genes, these display heterozygous sufficiency. We also generated F1 *Casp1*^−/−^*Casp11*^−/−^ mice by crossing *Casp1*^−/−^*Casp11*^−/−^ mice on B6 and 129 backgrounds. We assessed the ΔLD_100_ of four model pathogens (*L. monocytogenes, S*. Typhimurium, *B. thailandensis*, and *C. violaceum*) between WT and *Casp1*^−/−^*Casp11*^−/−^ mice on B6, 129, and F1 backgrounds. The results support the idea of Red Pawn barriers: caspase-1 and -11 protect against infection by ubiquitous environmental microbes but have minimal impact against host-adapted pathogens.

**Figure 1.**
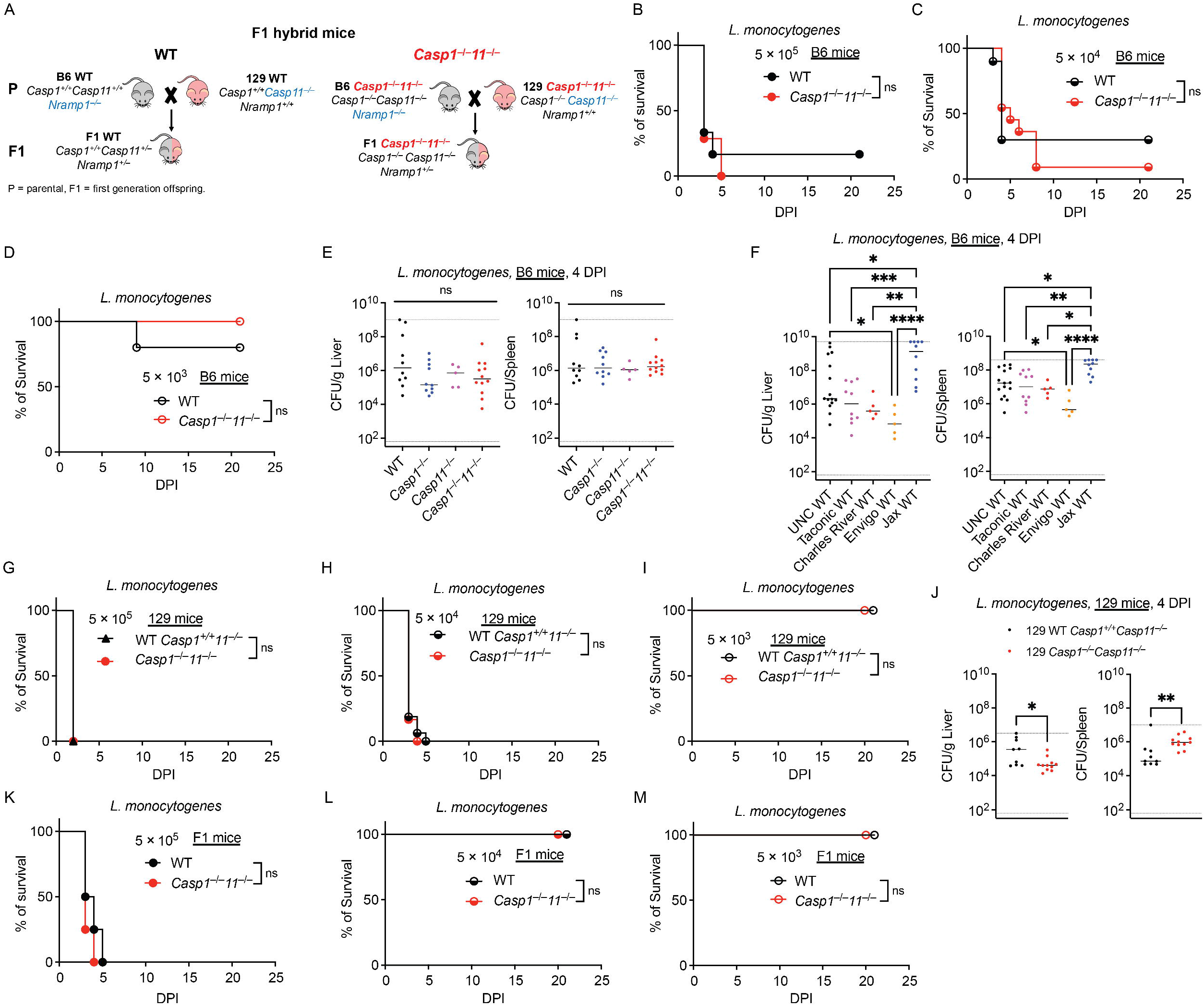
Caspase-1 and -11 are not required to control *L. monocytogenes* infection in B6, 129, and F1 mice. (A) F1 hybrid mice generation. C57BL/6 (B6) mice are *Nramp1*^mt/mt^ and 129 mice are *Casp11*^−/−^. F1 WT mice were generated by crossing B6 WT with 129 WT mice, the F1 WT mice were gene-sufficient for *Nramp1* and *Casp11* together with other unknown mutations in B6 and 129 mice. F1 *Casp1*^−/−^*Casp11*^−/−^ mice were generated by crossing B6 *Casp1*^−/−^*Casp11*^−/−^ with 129 *Casp1*^−/−^*Casp11*^−/−^ mice. F1 *Casp1*^−/−^*Casp11*^−/−^ mice were only deficient in caspase-1 and -11, but gene-sufficient for *Nramp1* together with other unknown mutations in B6 and 129 mice. Mice were infected intravenously (IV) with the indicated doses of *L. monocytogenes* (B–D, G–I, and K–M) or with 10^4^ CFU of *L. monocytogenes* (E, F, and J). Survival was monitored (B–D, G–I, and K–M), or mice were harvested at 4 DPI, and bacterial burdens in the liver and spleen were determined (E, F, and J). Mice: (B) WT=6, *Casp1*^−/−^*Casp11*^−/−^=7; (C) WT=10, *Casp1*^−/−^ *Casp11*^−/−^=11; (D) WT=5, *Casp1*^−/−^*Casp11*^−/−^=7; (E) WT=10, *Casp1*^−/−^=10, *Casp11*^−/−^=5, *Casp1*^−/−^ *Casp11*^−/−^=12; (F) UNC WT=15, Taconic WT=10, Charles River WT=5, Envigo WT=5, Jax WT=10; (G) WT=4, *Casp1*^−/−^*Casp11*^−/−^=5; (H) WT=16, *Casp1*^−/−^*Casp11*^−/−^=12; (I) WT=7, *Casp1*^−/−^ *Casp11*^−/−^=4; (J) WT=9, *Casp1*^−/−^*Casp11*^−/−^=11; (K) n=4 for both; (L) WT=4, *Casp1*^−/−^*Casp11*^−/−^=3; (M) n=3 for both. Data from B–E, G–I, and K–M are representative of at least two experiments. Data from F and J are pooled from two experiments. Bar represents median. Dashed lines indicate detection limit. Survival data were compared using the log-rank Mantel–Cox test. Bacterial burdens were analyzed using one-way ANOVA (E and F) or Student’s *t* test (J). ns, *p*□>□0.05; \**p*□<□0.05; \*\**p*□<□0.01; \*\*\**p*□<□0.001; \*\*\*\**p*□<□0.0001.

## Results

### Caspase-1 and -11 fail against Listeria monocytogenes

*L. monocytogenes* invades parenchymal cells using internalins and enters macrophages by phagocytosis; in both cases it escapes the vacuole using listeriolysin O and replicates in the host cytosol^33^. It is acquired from contaminated food and causes invasive disease mainly in young, elderly, immunocompromised, or pregnant individuals^34^, yet it also persists in diverse environmental reservoirs including water, soil, plants, and animals^35,36^. Thus, *L. monocytogenes* is a host-adapted pathogen that can also thrive in the environment. Cultured macrophages in vitro can detect *L. monocytogenes* through a variety of inflammasomes, including NLRC4, NLRP3, and AIM2^37–43^. However, studies in B6 mice show that this potential detection is not very efficacious in vivo. One study showed that whereas 80% of WT mice succumbed to 10^6^ CFU of *L. monocytogenes*, 100% of caspase-1/11-deficient mice succumbed^44^, suggesting that the protective effect of caspase-1 and -11 is only incremental. In contrast, another study showed that all B6 WT mice and *Asc*^−/−^ mice succumbed to 10^6^ CFU of *L. monocytogenes* infection, indicating no lethal dose change at this inoculum (ΔLD_100_ = 1), and surprisingly, *Asc*^−/−^ mice survived slightly longer than WT mice^45^. To more thoroughly determine the ΔLD_100_ in B6 mice, we infected B6 WT and B6 *Casp1*^−/−^*Casp11*^−/−^ mice with 5 × 10^5^, 5 × 10^4^, or 5 × 10^3^ CFU of *L. monocytogenes*. As in the previous publication using 10^6^ CFU infection, all or most B6 WT and B6 *Casp1*^−/−^*Casp11*^−/−^ mice died following 5 × 10^5^ and 5 × 10^4^ CFU inoculations with similar kinetics (Figures 1B and 1C). Conversely, at 5 × 10^3^ CFU, almost all mice of both genotypes survived infection (Figure 1D). The survival data from B6 mice indicated no lethal dose change between WT and caspase-1/11-deficient mice (ΔLD_100_ = 1). We then compared CFU burdens between B6 WT and caspase-1/11-deficient mice. Consistent with the survival data, WT mice had similar bacterial burdens at 4 DPI to those in *Casp1*^−/−^, *Casp11*^−/−^, and *Casp1*^−/−^*Casp11*^−/−^ mice (Figure 1E), suggesting that caspase-1 and -11 had no detectable role in defense against *L. monocytogenes* in B6 mice. These results contrast with several papers that show that caspase-1/11-deficient mice have higher *L. monocytogenes* burdens^44,46^. We wondered what could cause our inability to replicate these studies. Studies have shown that the microbiota affects the outcome of *L. monocytogenes* infection in mice^47,48^, which may explain discrepancies between studies showing that caspase-1 and -11 have a beneficial^44^ or detrimental role^45,49,50^. To validate the role of the microbiota during *L. monocytogenes* infection, we infected B6 WT mice from our own colony, and B6 WT mice ordered directly from four different vendors (the Jackson Laboratory, Taconic, Charles River Laboratories, and Envigo). We then determined the CFU burdens at 4 DPI. Surprisingly, B6 WT mice from diverse sources had a range of bacterial burdens in the liver and spleen, with mice from the Jackson Laboratory having remarkably higher bacterial burdens compared to B6 WT mice from the other four sources (Figure 1F), suggesting that the microbiota is an important factor to consider when interpreting in vivo *L. monocytogenes* infection data.

To test caspase-1 and -11 on an *Nramp1*^+/+^ background, we generated a 129 line in which both *Casp1* and the adjacent *Casp11* gene are deleted. We observed similar survival outcomes in WT and *Casp1*^−/−^*Casp11*^−/−^ mice on the 129 background. All 129 WT (*Casp1*^+/+^*Casp11*^−/−^) and 129 *Casp1*^−/−^*Casp11*^−/−^ mice succumbed to 5 × 10^5^ and 5 × 10^4^ CFU infection (Figures 1G and 1H), while 100% of them survived 5 × 10^3^ CFU infection (Figure 1I). Surprisingly, the bacterial burdens in the liver of 129 *Casp1*^−/−^*Casp11*^−/−^ mice at 4 DPI were slightly reduced compared to those in 129 WT mice, whereas the spleen burdens were slightly higher in 129 *Casp1*^−/−^*Casp11*^−/−^ mice (Figure 1J), suggesting that caspase-1 and -11 may have minor, tissue-specific effects. Nonetheless, the ΔLD_100_ in 129 mice was 1; caspase-1 and -11 had no effect on the survival of 129 mice during *L. monocytogenes* infection.

As in B6 and 129 mice, survival of F1 WT and F1 *Casp1*^−/−^*Casp11*^−/−^ mice did not differ significantly at any of the three inocula examined (Figures 1K–1M), indicating a ΔLD_100_ of 1 in F1 mice as well. Taken together, our data on B6, 129, and F1 mice strongly support the conclusion that caspase-1 and -11 are ineffective against *L. monocytogenes* in vivo.

### Caspase-1 and -11 fail against *Salmonella* Typhimurium

*S*. Typhimurium is a Gram-negative bacterium that encodes two T3SS, which are important for invading host cells and replicating within the *Salmonella*-containing vacuole^51^. Human infections with *S*. Typhimurium are common, with mild diarrhea and fever, and most people recover from infection without treatment^52^. *S*. Typhimurium is transmitted through the fecal-oral route, and the natural reservoirs of *S*. Typhimurium are vertebrates. Therefore, *S*. Typhimurium is a vertebrate-adapted pathogen. *S*. Typhimurium is a common model pathogen in the inflammasome field, although B6 WT mice succumb to systemic *S*. Typhimurium infection even at low infection doses due in large part to the *Nramp1* mutation^53^. In prior publications, investigators have infected WT and *Casp1*^−/−^*Casp11*^−/−^ mice on the B6 background with 10^5^ to 10^8^ CFU of *S*. Typhimurium orally; all mice succumbed to infection, although WT mice succumbed 2–3 days later^5,6^. Thus, orally infected B6 mice did not have a ΔLD_100_ within inocula from 10^5^ to 10^8^ CFU. After systemic infection (i.e., IP infection), B6 mice are very susceptible to *S*. Typhimurium, with lethal doses of less than 10 CFU; at this minimal dose, B6 *Casp1*^−/−^ mice succumbed with similar kinetics (Figures 2A–2C). This extreme susceptibility makes it impossible to assess ΔLD_100_ in B6 mice.

**Figure 2.**
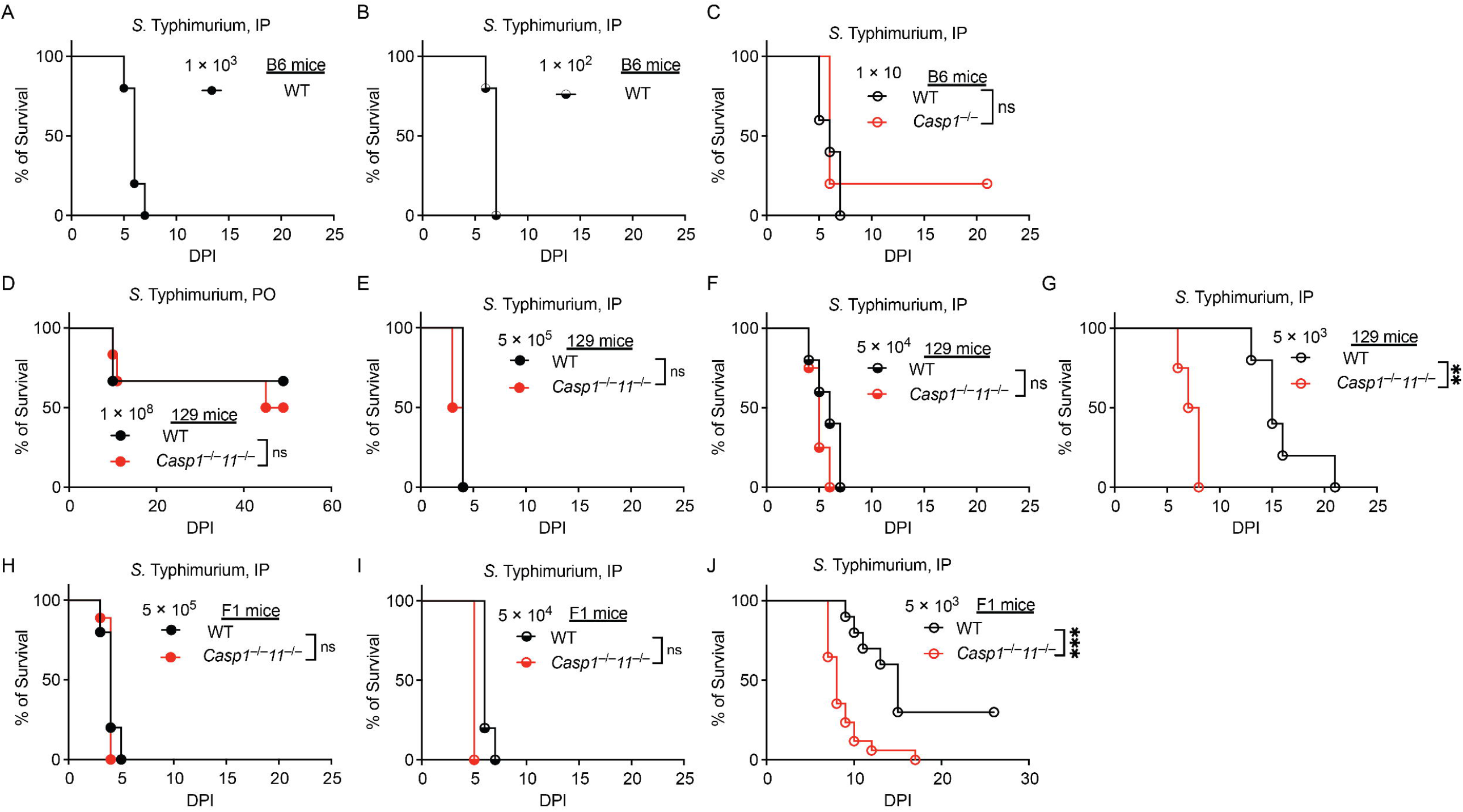
Caspase-1 and -11 are not essential for controlling *S*. Typhimurium infection in B6, 129, and F1 mice. Mice were infected intraperitoneally (IP) with the indicated doses of *S*. Typhimurium, except in (D), where mice were infected orally. Survival was monitored. Mice: (A– C) n=5 for all; (D) n=6 for both; (E) n=4 for both; (F and G) WT=5, *Casp1*^−/−^*Casp11*^−/−^=4; (H) WT=5, *Casp1*^−/−^*Casp11*^−/−^=9; (I) WT=5, *Casp1*^−/−^*Casp11*^−/−^=7; (J) WT=10, *Casp1*^−/−^*Casp11*^−/−^=17. Data are pooled from two experiments (A–C), or representative of at least two experiments (D– J). Survival data were compared using the log-rank Mantel–Cox test. ns: not significant; * *p*l< 0.05; \*\**p*□<□0.01; *** *p*l< 0.001.

We therefore turned to the 129 mice infection model because these mice have an intact *Nramp1* gene and are more resistant to *S*. Typhimurium. Orally infected 129 mice did not have a ΔLD_100_ at an inoculum of 10^8^ CFU (Figure 2D). During systemic infection there was no significant difference in survival between 129 WT and 129 *Casp1*^−/−^*Casp11*^−/−^ mice inoculated with 5 × 10^5^ and 5 × 10^4^ CFU, as all mice died at similar times (Figures 2E and 2F). At the lowest inoculum dose tested (5 × 10^3^ CFU), 129 WT mice survived significantly longer (> 7 days) than 129 *Casp1*^−/−^*Casp11*^−/−^ mice, but all mice still succumbed to infection (Figure 2G), indicating no lethal dose change (ΔLD_100_ = 1).

To assess the importance of caspase-1 and -11 in a genetically complex background, we next examined F1 mice. All F1 WT and F1 *Casp1*^−/−^*Casp11*^−/−^ mice succumbed to systemic infection with inocula of 5 × 10^5^ and 5 × 10^4^ CFU with similar kinetics (Figures 2H and 2I). When inoculated with 5 × 10^3^ CFU, F1 WT mice survived significantly longer than F1 *Casp1*^−/−^*Casp11*^−/−^ mice, and 30% of F1 WT mice survived infection, whereas 100% of F1 *Casp1*^−/−^*Casp11*^−/−^ mice succumbed to infection (Figure 2J), indicating that caspase-1 and -11 play a role at this lower inoculum in F1 mice. In conclusion, data from 129 and F1 mice suggested that caspase-1 and - 11 fail against *Salmonella* Typhimurium when the inocula were higher than 5 × 10^4^ CFU, whereas at a lower inoculum (5 × 10^3^ CFU), caspase-1 and -11 may delay the infection kinetics, but do not rescue most mice. We therefore estimate there is a ΔLD_100_ of greater than 1. Broken down by background, the ΔLD_100_ could not be determined in B6 mice, which succumb to fewer than 10 CFU regardless of genotype; it was 1 in 129 mice; and it was greater than 1 in F1 mice, where 30% of WT mice survived the lowest inoculum while all caspase-1/11-deficient mice succumbed.

### Caspase-1 and -11 defend against *Burkholderia thailandensis*

*B. thailandensis* is a Gram-negative bacterium that encodes a T3SS similar to that of *Shigella*. After being taken up by host cells through T3SS-mediated invasion or phagocytosis, *B. thailandensis* uses its T3SS to escape the phagosome and replicate in the host cytosol^27^. *B. thailandensis* is ubiquitous in soil in tropical regions^54^ but its natural host is unknown. Human infection by *B. thailandensis* is exceedingly rare, with only three case reports to date^55–57^. Our laboratory previously studied *B. thailandensis* on the B6 mouse background. B6 WT mice were resistant to 6 × 10^7^ CFU of *B. thailandensis*, while B6 *Casp1*^−/−^*Casp11*^−/−^ mice succumbed to 100 CFU^28^. We verified our previous finding in a new cohort of mice (Figures 3A–3C).

**Figure 3.**
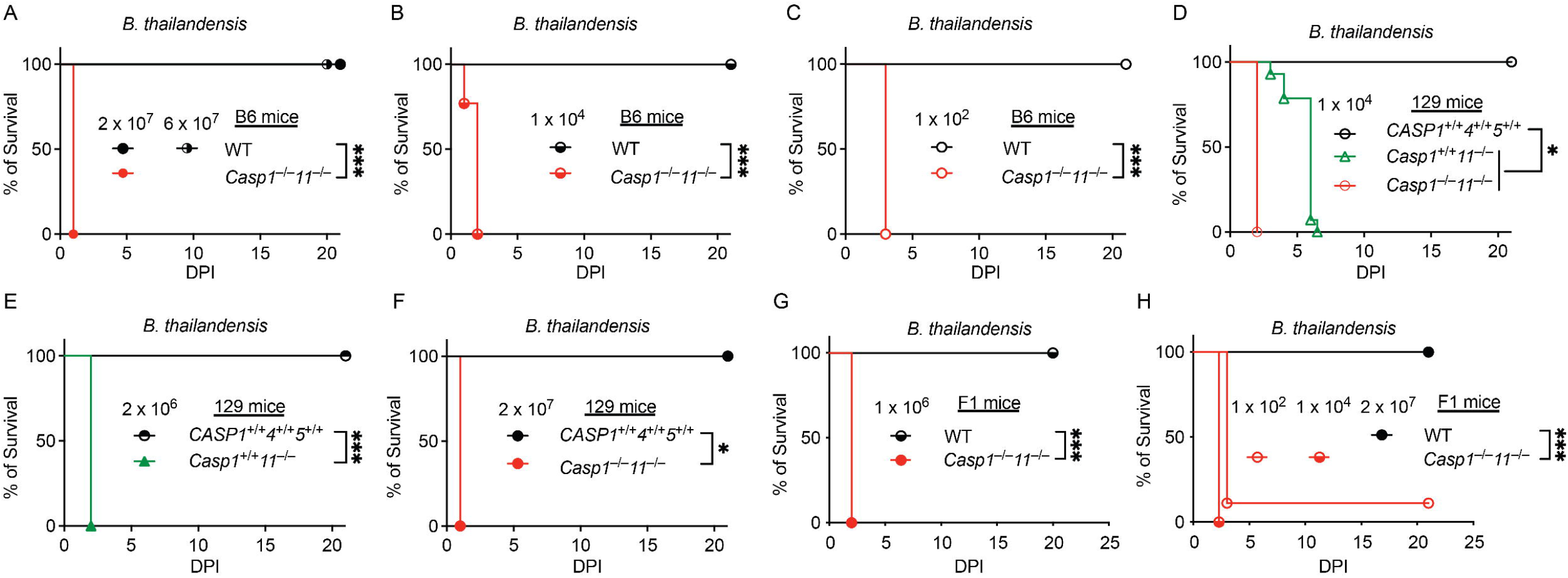
Caspase-1 and -11 are required for defense against *B. thailandensis* infection in B6, 129, and F1 mice. Mice were infected intraperitoneally (IP) with the indicated doses of *B. thailandensis*. Survival was monitored. Mice: (A) WT=5 for both doses, *Casp1*^−/−^*Casp11*^−/−^=6; (B) WT=5, *Casp1*^−/−^*Casp11*^−/−^=13; (C) WT=5, *Casp1*^−/−^*Casp11*^−/−^=11; (D) *CASP1*^+/+^*CASP4*^+/+^*CASP5*^+/+^=3; *Casp1*^+/+^*Casp11*^−/−^=14, *Casp1*^−/−^*Casp11*^−/−^=3; (E) *CASP1*^+/+^*CASP4*^+/+^*CASP5*^+/+^=6; *Casp1*^+/+^*Casp11*^−/−^=10; (F) *CASP1*^+/+^*CASP4*^+/+^*CASP5*^+/+^=2; *Casp1*^−/−^*Casp11*^−/−^=5; (G) WT=5, *Casp1*^−/−^*Casp11*^−/−^=8; (H) WT=9, *Casp1*^−/−^*Casp11*^−/−^ 1 × 10^4^=8; *Casp1*^−/−^*Casp11*^−/−^ 1 × 10^2^=9. Data are representative of at least two experiments. Data were compared using the log-rank Mantel–Cox test. * *p*l< 0.05; *** *p*l< 0.001.

Turning to the 129 background, we hypothesized that the endogenous *Casp11* mutation would cause susceptibility to *B. thailandensis* infection even in the presence of a functional *Nramp1* gene. Indeed, 129 WT (*Casp1*^+/+^*Casp11*^−/−^) mice were susceptible to *B. thailandensis* infection (Figures 3D and 3E). Mice with an additional deletion of *Casp1* on the 129 background (*Casp1*^−/−^*Casp11*^−/−^) also died acutely from infection, but faster than the *Casp1*-sufficient mice (Figure 3D). We also engineered 129 WT mice to express the human version of the caspase locus, replacing the murine 129 *Casp1*^+/+^*Casp11*^−/−^ locus with the human *CASP1*^+/+^*CASP4*^+/+^*CASP5*^+/+^ locus; this restores cytosolic LPS sensing in 129 mice (Koller et al., manuscript in preparation). These mice were resistant to infection with up to 2 × 10^7^ CFU (Figures 3E and 3F), indicating that human CASP1, CASP4, and CASP5 can clear *B. thailandensis* in a 129 background. We conclude that the endogenous *Casp11* mutation, and no other uncharacterized mutations, causes susceptibility in 129 WT mice.

We hypothesized that the susceptibility of 129 WT mice would be rescued by crossing with B6 mice to obtain F1 mice. Indeed, F1 WT mice were fully resistant to *B. thailandensis* infection, even at high doses of 2 × 10^7^ CFU (Figures 3G and 3H). In contrast, F1 *Casp1*^−/−^*Casp11*^−/−^ mice succumbed to 100 CFU infection (Figures 3G and 3H). Therefore, caspase-1 and -11 provide potent defenses against *B. thailandensis* in B6, 129, and F1 mice. Based on the doses tested, the ΔLD_100_ was at least 6 × 10^5^ in B6 mice (WT resistant to 6 × 10^7^ CFU versus 10^2^ CFU in caspase-1/11-deficient mice) and at least 2 × 10^5^ in F1 mice (2 × 10^7^ versus 10^2^ CFU); in 129 mice the ΔLD_100_ was 1, because 129 WT mice are themselves caspase-11-deficient.

### Caspase-1 and -11 defend against *Chromobacterium violaceum*

*C. violaceum* is a Gram-negative bacterium encoding a T3SS, which is like the SPI1 T3SS of *Salmonella*^58^. *C. violaceum* is ubiquitous in soil and freshwater^59^. *C. violaceum* rarely infects immunocompetent individuals, and human case reports of *C. violaceum* infection are usually seen in immunocompromised patients or near-drowning victims^59^. Our laboratory has found that caspase-1 and -11 are essential to clear *C. violaceum* infection in B6 mice^29^, which was reproduced here (Figures 4A and 4B). We calculate the ΔLD_100_ as > 1,000.

**Figure 4.**
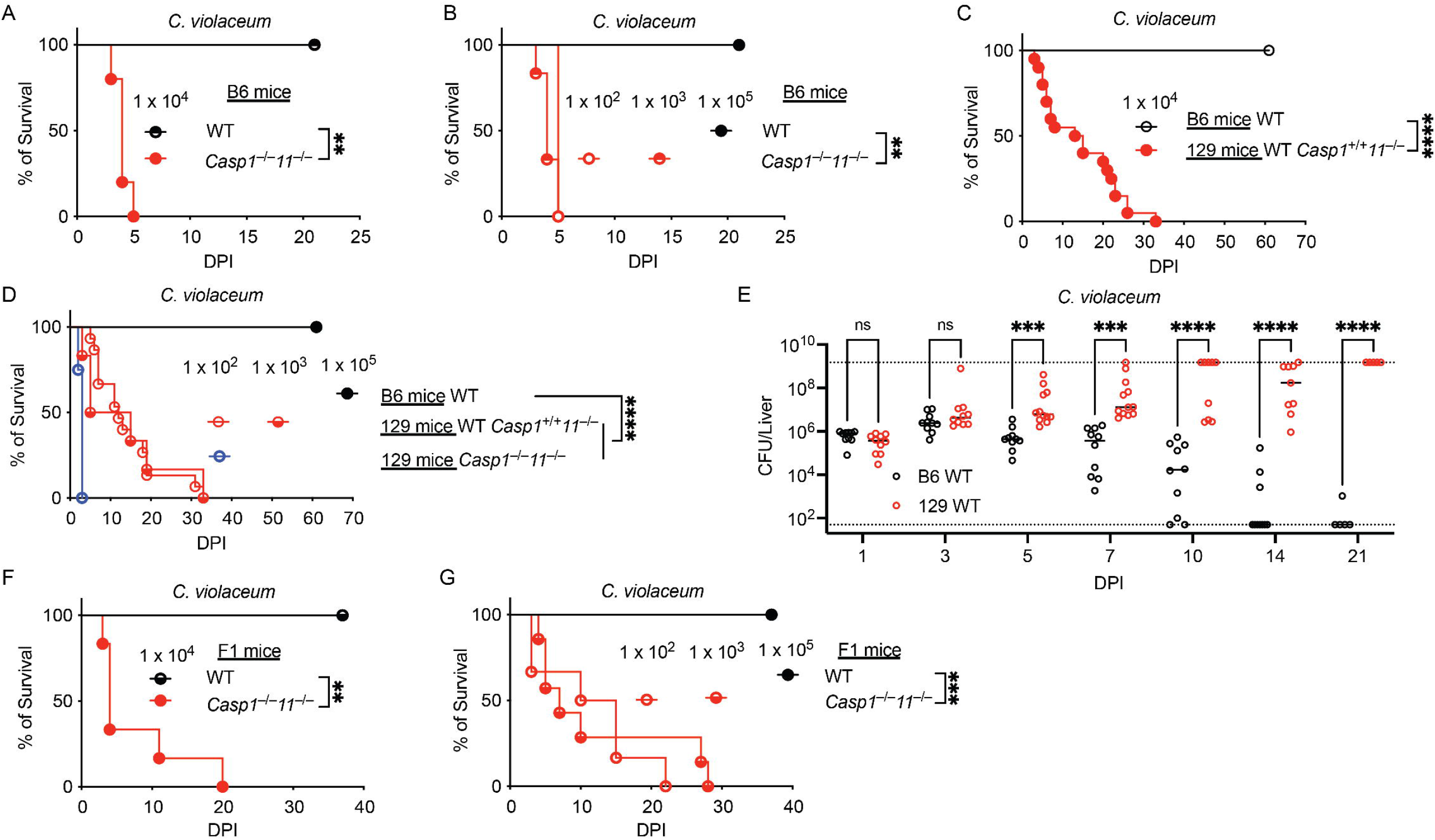
Caspase-1 and -11 are required to control *C. violaceum* infection in B6, 129, and F1 mice. Mice were infected intraperitoneally (IP) with the indicated doses of *C. violaceum* (A–D, F, and G) or with 10^4^ CFU of *C. violaceum* (E). Survival was monitored (A–D, F, and G), or mice were harvested at the indicated times post-infection, and bacterial burdens in the liver and spleen were determined (E). Mice: (A) n=5 for both; (B) WT=5, *Casp1*^−/−^*Casp11*^−/−^ 1 × 10^3^=6, *Casp1*^−/−^*Casp11*^−/−^ 1 × 10^2^=5; (C) B6 WT=9, 129 WT=20; (D) B6 WT=11, 129 WT 1 × 10^3^=6, 129 WT 1 × 10^2^=15, 129 *Casp1*^−/−^*Casp11*^−/−^=4; (E) n=10 for both at 1, 3, 10, and 14 DPI, B6 WT=10 for 5 and 7 DPI, 129 WT=12 for 5 and 7 DPI, n=5 for both at 21 DPI; (F) WT=4, *Casp1*^−/−^ *Casp11*^−/−^=6; (G) WT=7, *Casp1*^−/−^*Casp11*^−/−^ 1 × 10^3^=7, *Casp1*^−/−^*Casp11*^−/−^ 1 × 10^2^=6. Data from A and B are representative of at least two experiments. Data from C–G are pooled from two experiments. Bar represents median. Dashed lines indicate detection limit. Survival data were compared using the log-rank Mantel–Cox test. Bacterial burdens were analyzed using two-way ANOVA. ns, *p*□>□0.05; \*\**p*□<□0.01; \*\*\**p*□<□0.001; \*\*\*\**p*□<□0.0001.

Turning to the 129 background, surprisingly, 100% of 129 WT (*Casp1*^+/+^*Casp11*^−/−^) mice succumbed to *C. violaceum* infection, even at low doses of 100 CFU (Figures 4C and 4D), with increasing bacterial burdens over time (Figure 4E). A role of caspase-1 and -11 was still present because 129 *Casp1*^−/−^*Casp11*^−/−^ mice succumbed significantly faster than 129 WT mice; however, in the 129 background, the ΔLD_100_ = 1.

The susceptibility of caspase-1-sufficient 129 WT mice indicates that additional susceptible loci are present in 129 WT mice. We hypothesized that these would be rescued by crossing with B6 mice to obtain genetically complex F1 backgrounds. Indeed, F1 mice were highly resistant to *C. violaceum* infection (Figures 4F and 4G). In contrast, F1 *Casp1*^−/−^*Casp11*^−/−^ mice succumbed to as few as 100 CFU infection (Figures 4F and 4G). Therefore, caspase-1 and -11 protect against lethal *C. violaceum* in B6 and F1 mice, but other loci cause lethality in 129 mice.

## Discussion

Under the idea of Red Pawn barriers, ubiquitous environmental pathogens (personified as Red Pawns) are not in constant evolution with their non-hosts (e.g., humans and mice), although they carry virulence traits (e.g., T3SS and T6SS) against natural hosts that presumably do not encode caspase-1 and -11. Thus, there is no selective pressure for Red Pawns to develop caspase-1 and -11 evasion strategies, and they can be readily cleared by caspase-1 and -11.

We determined the importance of caspase-1 and -11 in defense against two host-adapted pathogens (*L. monocytogenes* and *S*. Typhimurium) and two environmental pathogens (*B. thailandensis* and *C. violaceum*) in WT and *Casp1*^−/−^*Casp11*^−/−^ mice on B6, 129, and F1 backgrounds. Our results showed that caspase-1 and -11 fail against host-adapted pathogens but successfully control environmental pathogens. Our results support the idea that caspase-1 and -11 achieved a near-permanent victory against environmental pathogens, forming Red Pawn barriers that prevent these organisms from engaging in the Red Queen’s race with mice and likely humans. In contrast, host-adapted pathogens evade caspase-1 and -11 defenses.

Our conclusions rest on the fold change in lethal dose rather than on burdens or survival at a single inoculum. Protection that looks absolute at one dose can shrink considerably once the dose is titrated. Our own work with *Burkholderia pseudomallei* is the cautionary case: at a dose chosen (100 CFU) because it was sublethal in WT mice, all *Casp1*^−/−^*Casp11*^−/−^ mice died while all WT mice survived, which appeared to be complete protection^15,60^. Re and colleagues tested several doses: 25 CFU of *B. pseudomallei* killed *Casp1*^−/−^ mice while WT mice survived, and 200 CFU killed every WT mouse, placing the true shift closer to 8-fold^61^. Most published studies report a single inoculum (Table 1) and therefore cannot distinguish an 8-fold barrier from a 10^6^-fold one. Titrating the challenge dose of four pathogens allows us to determine the magnitude of protection thoroughly. The separations we observe are not incremental.

These data also bear on the alternative proposal that caspase-1 and -11 are maintained because they delay infection rather than prevent it, buying the host time to mount an adaptive response. The delay is real: 129 and F1 WT mice survived significantly longer than *Casp1*^−/−^ *Casp11*^−/−^ mice at the lowest *S*. Typhimurium inoculum, and mice that survive *L. monocytogenes* infection to 7–8 days mount a cytotoxic T lymphocyte response sufficient to clear the bacterium^62^. For both host-adapted pathogens, however, the delay does not become a change in lethal dose. Against *B. thailandensis* and *C. violaceum*, the same genotypes shifted the lethal dose by up to six orders of magnitude. A selective pressure that bought only a few days would not produce a gap of this size between the two classes of pathogen.

Being environmental does not by itself make a microbe a Red Pawn of caspase-1 and -11. *Klebsiella pneumoniae* and *Vibrio vulnificus* cause disease mainly in compromised hosts, yet neither encodes a T3SS nor invades the cytosol, and neither produces more than a marginal phenotype in caspase-1/11-deficient mice^15^. It is likely that other Red Pawn barriers hold these environmental pathogens at bay. Indeed, mice and people deficient in the phagocyte NADPH oxidase succumb to even 4 × 10^3^ CFU of the environmental bacterial pathogen *Burkholderia cepacia*^63^, whereas WT mice survive 10^7^ CFU infection. Caspase-1 and -11 are not important for defense against a patient isolate of *Burkholderia cepacia*^29^. Therefore, caspase-1 and -11 can only defend against bacteria they can sense. *B. thailandensis* and *C. violaceum* are sensed because they use a T3SS and invade the cytosol. For the same reason we prefer the term environmental pathogen to opportunistic pathogen, since infection of an immunocompromised host is an evolutionary dead end that exerts no selective pressure on the microbe (Miao, 2026, under review).

The durability of this barrier follows from what the sensors detect. Rod, needle and flagellin subunits reach the cytosol as accidental byproducts of T3SS injection, and LPS becomes visible the moment a bacterium enters the cytosol; escaping detection therefore requires either abandoning the intracellular niche or perfecting an intrinsically imperfect injection apparatus (Miao, 2026, under review). Host-adapted pathogens have taken the first route, but only under the sustained selection that accompanies host-to-host transmission. Environmental microbes never experience that pressure, so a barrier that would eventually become porous to a transmitting pathogen holds against them indefinitely.

The barrier is also layered rather than singular. 129 WT mice are caspase-1-sufficient yet succumbed to 10^2^ CFU of *C. violaceum*, indicating that additional loci contribute to clearance in this strain, and patients with chronic granulomatous disease are susceptible to *C. violaceum* despite intact caspase-1 and -11^64^, placing the NADPH oxidase and pyroptosis as Red Pawn barriers that act in series to prevent *C. violaceum* infection. A ΔLD_100_ measured in any one inbred strain therefore reports the combined state of several loci. This is precisely why we moved to genetically complex F1 hybrid mice.

As already discussed in a review article our group published^15^, the idea of Red Pawn barriers has a few caveats. First, species barriers between mice and humans sometimes determine pathogen virulence. For example, *Francisella novicida* is an environmental pathogen associated with brackish water^65^. Human infections with *F. novicida* are extremely rare^66^. However, B6 WT mice are extremely susceptible to systemic *F. novicida* infection, with a lethal dose lower than 10 CFU after systemic infection^67^, suggesting that Red Pawn barriers exist between mice and humans. Indeed, even within the inflammasome, caspase-1, and caspase-11 defense pathways there are numerous differences between mice and humans. For example, mice encode a single *Casp11* whereas humans have duplicated the gene into *CASP4* and *CASP5*. Another caveat of our approach is that we focus on systemic infection. We chose this route because it provides an advantage to the pathogen, allowing it to express its virulence maximally, and it minimizes the effects of microbiota and is a simplified model that can be compared across different pathogen types. For example, it is impossible to compare oral challenge routes for a gastrointestinal pathogen to subcutaneous infection routes for a skin pathogen.

Although we have found two Red Pawns held at bay by caspase-1 and -11, it is still surprising to us that a single innate immune gene deficiency would make normal-appearing mice susceptible to infection with 100 or fewer environmental bacteria, considering the fail-safe backup pathways built into our innate immune system^26^. It is likely that more environmental pathogens are defended against redundantly by our innate immune defenses, and identifying them could lead to new discoveries in innate immune defenses.

## Materials and Methods

### Mice

The following mouse lines were housed in a specific-pathogen-free facility at the University of North Carolina (UNC) at Chapel Hill or Duke University: wild-type C57BL/6 (Jackson # 000664), C57BL/6 *Casp1*^−/−^*Casp11*^−/−^, 129 WT (129S6/SvEv), 129 *Casp1*^−/−^*Casp11*^−/−^ and 129 *CASP1*^+/+^*CASP4*^+/+^*CASP5*^+/+^. The generation and characterization of 129 *Casp1*^−/−^*Casp11*^−/−^ and 129 *CASP1*^+/+^*CASP4*^+/+^*CASP5*^+/+^ will be described in more detail in a separate manuscript in preparation by Dr. Koller. All animal experiments used 8- to 12-week-old mice. Both males and females were included. Animal protocols were approved by the Institutional Animal Care and Use Committee (IACUC) at UNC-Chapel Hill or Duke University and met the US National Institutes of Health guidelines.

### Bacteria

*Listeria monocytogenes* (10403s), *S. enterica* serovar Typhimurium (SL1344), *Burkholderia thailandensis* (E264), and *Chromobacterium violaceum* (ATCC 12472) have been described previously^8,68^. *L. monocytogenes* and *C. violaceum* were grown in BHI. *S*. Typhimurium and *B. thailandensis* were grown in LB.

### In vivo infection

*L. monocytogenes* diluted in 200 μL of PBS was administered to mice by intravenous (IV) injection using a 30G needle. *S*. Typhimurium, *C. violaceum*, and *B. thailandensis* diluted in 200 μL of PBS were administered to mice by intraperitoneal (IP) injection using a 30G needle.

### CFU determination

After euthanasia, livers and spleens were removed from the mice and transferred to 1 mL of PBS. Organs were weighed and homogenized. Serial dilutions (1:10) were performed in 96-well plates. 40 μL of each dilution (4 × 10 μL) was plated on BHI or LB agar plates and incubated overnight at 37 °C.

### Statistics

All statistical analyses were performed using GraphPad Prism (version 11.0.2). Statistical significance was determined using an unpaired two-tailed Student’s *t* test for comparisons between two groups, one-way ANOVA with Tukey’s multiple-comparisons test for comparisons among three or more groups, two-way ANOVA with Tukey’s multiple-comparisons test for experiments involving two independent variables, or the log-rank (Mantel–Cox) test for survival analyses. Statistical significance was defined as follows: ns, *p*□>□0.05; \**p*□<□0.05; \*\**p*□<□0.01; \*\*\**p*□<□0.001; \*\*\*\**p*□<□0.0001.

## Acknowledgments

This work was supported by NIH grants AI136920, AI175078, AI181815, and AI198309 (E.A.M.).

